# PeakPerformance - a tool for Bayesian inference-based fitting of LC-MS/MS peaks

**DOI:** 10.1101/2024.02.17.580815

**Authors:** Jochen Nießer, Michael Osthege, Eric von Lieres, Wolfgang Wiechert, Stephan Noack

## Abstract

A major bottleneck of chromatography-based analytics has been the elusive fully automated identification and integration of peak data without the need of extensive human supervision. The presented Python package `PeakPerformance` applies Bayesian inference to chromatographic peak fitting, and provides an automated approach featuring model selection and uncertainty quantification. Currently, its application is focused on data from targeted liquid chromatography tandem mass spectrometry (LC-MS/MS), but its design allows for an expansion to other chromatographic techniques. `PeakPerformance` is implemented in Python and the source code is available on GitHub. It is unit-tested on Linux and Windows and accompanied by general introductory documentation, as well as example notebooks.

**Practical application:** The presented `PeakPerformance` tool performs automated chromatographic peak data fitting using Bayesian methodology. Accordingly, it innovates by delivering built-in uncertainty quantification for each peak, thus taking the measurement noise into account. Using a convergence statistic and based on the determined peak uncertainties, the differentiation of signals into peak and noise was improved and false positives or negatives were largely eliminated. The provided documentation and the implemented convenience functions are meant to lower the barrier of entry for users with little programming experience. Lastly, the modular design of the software enables modification and expansion to data from different chromatographic methods.

## 1 Introduction

In biotechnological research and industrial applications, chromatographic techniques are ubiquitously used to analyze samples from fermentations, e.g. to determine the concentration of substrates and products. Over the course of a regular lab-scale bioreactor fermentation, dozens to hundreds of samples and subsequently – depending on the number of analytes per sample – hundreds to thousands of chromatographic peaks may accrue. This is exacerbated by the spread of microbioreactors causing a further increase in the amount of samples per time [1, 2]. While the recognition and integration of peaks by vendor software is – in theory – automated, it typically requires visual inspection and occasional manual re-integration by the user due to a large number of false positives, false negatives or incorrectly determined baselines, ultimately downgrading it to a semi-automated process. Since this is a time-consuming, not to mention tedious, procedure and introduces the problem of comparability between purely manual and algorithm-based integration as well as user-specific differences, we instead propose a peak fitting solution based on Bayesian inference.

The advantage of this approach is the complete integration of all relevant parameters – i.e. baseline, peak area and height, mean, signal-to-noise ratio etc. – into one single model through which all parameters are estimated simultaneously. Furthermore, Bayesian inference comes with uncertainty quantification for all peak model parameters, and thus does not merely yield a point estimate as would commonly be the case. It also grants access to novel metrics for avoiding false positives and negatives by rejecting signals where a) a convergence criterion of the peak fitting procedure was not fulfilled or b) the uncertainty of the estimated parameters exceeded a user-defined threshold.

## 2. Materials and Methods

### 2.1 Implementation

`PeakPerformance` is an open source Python package compatible with Windows and Linux/Unix platforms. At the time of manuscript submission, it features three modules: pipeline, models, and plotting. Due to its modular design, `PeakPerformance` can easily be expanded by adding e.g. additional models for deviating peak shapes or different plots. Currently, the featured peak models describe peaks in the shape of normal or skew normal distributions, as well as double peaks of normal or skewed normal shape. The normal distribution is regarded as the ideal peak shape and common phenomena like tailing and fronting can be expressed by the skew normal distribution [3]. A third distribution commonly used to describe peak shapes is the exponentially modified Gaussian [4] for which an implementation in `PeakPerformance` is pending.

Bayesian inference is conducted utilizing the PyMC package [5] with the external sampler `nutpie` for improved performance [6]. Both model selection and analysis of inference data objects are realized with the ArviZ package [7]. Since the inference data is stored alongside graphs and report sheets, users may employ the ArviZ package or others for further analysis of the results if necessary.

### 2.2 Validation of PeakPerformance

Several stages of validation were employed to prove the suitability of `PeakPerformance` for chromatographic peak data analysis. The goals were to showcase the efficacy of `PeakPerformance` utilizing noisy synthetic data, investigate cases where a peak could reasonably be fit with either of the single peak models, and finally use experimental data to compare results obtained with `PeakPerformance` to those from the commercial vendor software Sciex MultiQuant.

For the first test, 500 random data sets were generated with the NumPy random module by drawing from the normal distributions detailed in Table 1 except for the mean parameter which was held constant at a value of 6. Subsequently, normally distributed random noise (𝒩(0, 0.6) or 𝒩(0, 1.2) for data sets with the tag “higher noise”) was added to each data point. The amount of data points per time was chosen based on an LC-MS/MS method routinely utilized by the authors and accordingly set to one data point per 1.8 s.

**Table 1:** Normal distributions from which parameters were drawn randomly to create synthetic data sets for the validation of PeakPerformance.

| parameter | model |  |
| --- | --- | --- |
|  | 1<sup>st</sup> test | 2<sup>nd</sup> test |
| area | $\mathcal{N}(8, 0.5)$ | - |
| standard deviation | $\mathcal{N}(0.5, 0.1)$ | $\mathcal{N}(0.5, 0.1)$ |
| skewness | $\mathcal{N}(0, 2)$ | - |
| baseline intercept | $\mathcal{N}(25, 1)$ | $\mathcal{N}(25, 1)$ |
| baseline slope | $\mathcal{N}(0, 1)$ | $\mathcal{N}(0, 1)$ |

In marginal cases when the shape of a single peak had a slight skew, the automated model selection would at times settle on a normal or a skew normal model. Therefore, it was relevant to investigate whether this choice would lead to a significant discrepancy in estimated peak parameters. Accordingly, for the second test synthetic data sets were generated with the NumPy random module according to Table 1 and noise was added as described before. The residual parameters were held constant, i.e. the mean was fixed to 6, the area to 8, and the skewness parameter *α* to 1.

For the third and final test, experimental peak data was analyzed with both `PeakPerformance` (version 0.7.0) and Sciex MultiQuant (version 3.0.3) with human supervision, i.e. the results were visually inspected and corrected if necessary. The data set consisted of 192 signals comprised of 123 single peaks, 50 peaks as part of double peaks, and 19 noise signals.

### 2.3 Composition and assumptions of peak models

Peak models in `PeakPerformance` require the definition of prior probability distributions (priors) for their parameters as well as the choice of an intensity function and a likelihood function. Generally, priors are derived from a given time series and given a weakly informative parametrization, such that the resulting inferences of parameters like the peak height are primarily based on the data. While defining priors in a data-dependent manner is generally to be avoided, it is clearly not tenable to define legitimate priors for all kinds of different peaks with heights and areas varying by multiple orders of magnitude and retention times, i.e. mean values, scattered across the whole run time of the LC-MS/MS method. In order to flexibly build models for all these peaks in an automated manner and embedded in a standardized data pipeline, some parameter priors had to be based on the raw data. If specific distributions or their parameters had to be restricted to certain value ranges, error handling was incorporated. For example, when only positive values were acceptable or when 0 was not a permissive value, a lower bound was defined using NumPy’s clip function.

Regarding shared model elements across all intensity functions, one such component of all models presented hereafter is the likelihood function

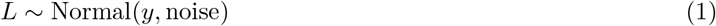

with *y* as the predicted intensity and noise as the free parameter describing the standard deviation of measurement noise. This definition encodes the assumption that observed intensities are the result of normally distributed noise around the true intensity values of a peak. In turn, the noise parameter is defined as

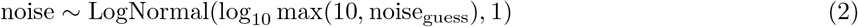

The log-normal distribution where the logarithm of the random variable follows a normal distribution was chosen partly to exclude negative values from the solution space and also due to its shape attributing a higher fraction of the probability density to lower values provided the standard deviation is defined sufficiently high. This prior is defined in a raw data-dependent manner as the noise_guess_ amounts to the standard deviation of the differences of the first and final 15 % of intensity values included in a given time frame and their respective mean values.

The intensity function itself is defined as the sum of a linear baseline function and a peak intensity function, the latter of which is composed of a given distribution’s probability density function (PDF) scaled up to the peak size by the area or height parameter. The linear baseline

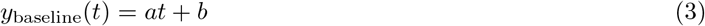

features the slope and intersect parameters *a* and *b*, respectively, both of which were given a normally distributed prior. The data-dependent guesses for these priors are obtained by constructing a line through the means of the first and last three data points of a given intensity data set which oftentimes already resulted in a good fit. Hence, the determined values for slope (*a*_guess_) and intercept (*b*_guess_) are used as the means for their pertaining priors and the standard deviations are defined as fractions of them with minima set to 0.5 and 0.05, respectively. Here, the exact definition of the standard deviations was less important than simply obtaining an uninformative prior which, while based on the rough fit for the baseline, possesses a sufficient degree of independence from it, thus allowing deviations by the Bayesian parameter estimation.

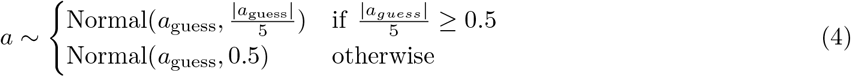

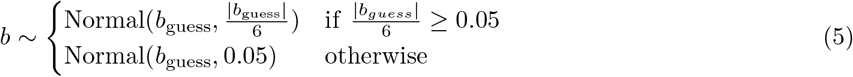

The initial guesses noise_guess_, *a*_guess_, and *b*_guess_ are calculated from raw time and intensity by the `initial guesses()` function from the `models` submodule.

Beyond this point, it is sensible to categorize models into single and double peak models since these subgroups share a larger common basis. Starting with single peak models, the normal-shaped model (Figure 1a) requires only three additional parameters for defining its intensity function. The mean value *µ* has a normally distributed prior with the center of the selected time frame min 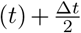 as its mean and 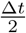 as the standard deviation where Δ*t* corresponds to the length of the time frame. Accordingly, the resulting prior is rather compressed and weakly informative. The prior for the standard deviation of the normal-shaped peak model was defined with a half-normal distribution, once again to avoid values equaling or below 0. As a half normal distribution only features a standard deviation, this was set to 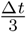. The final parameter is the peak height used for scaling up the distribution to match the size of the peak. Here, a rather uninformative half-normal distribution with a scale amounting to 95 % of the highest intensity value in the time frame was selected The second featured single peak model is based on the skew normal distribution (Figure 1b) which has an additional skewness parameter *α* enabling a one-sided distortion of the peak or resulting in identity to the normal-shaped peak model when *α* = 0. Hence, the prior of *α* is constituted by a normal distribution centered on 0 with a standard deviation of 3.5 to allow for a sufficiently large range of possible values for *α* and thus a realistic skew. Instead of the peak height, the peak area was utilized to scale the distribution, albeit with an identical prior.

**Figure 1:**
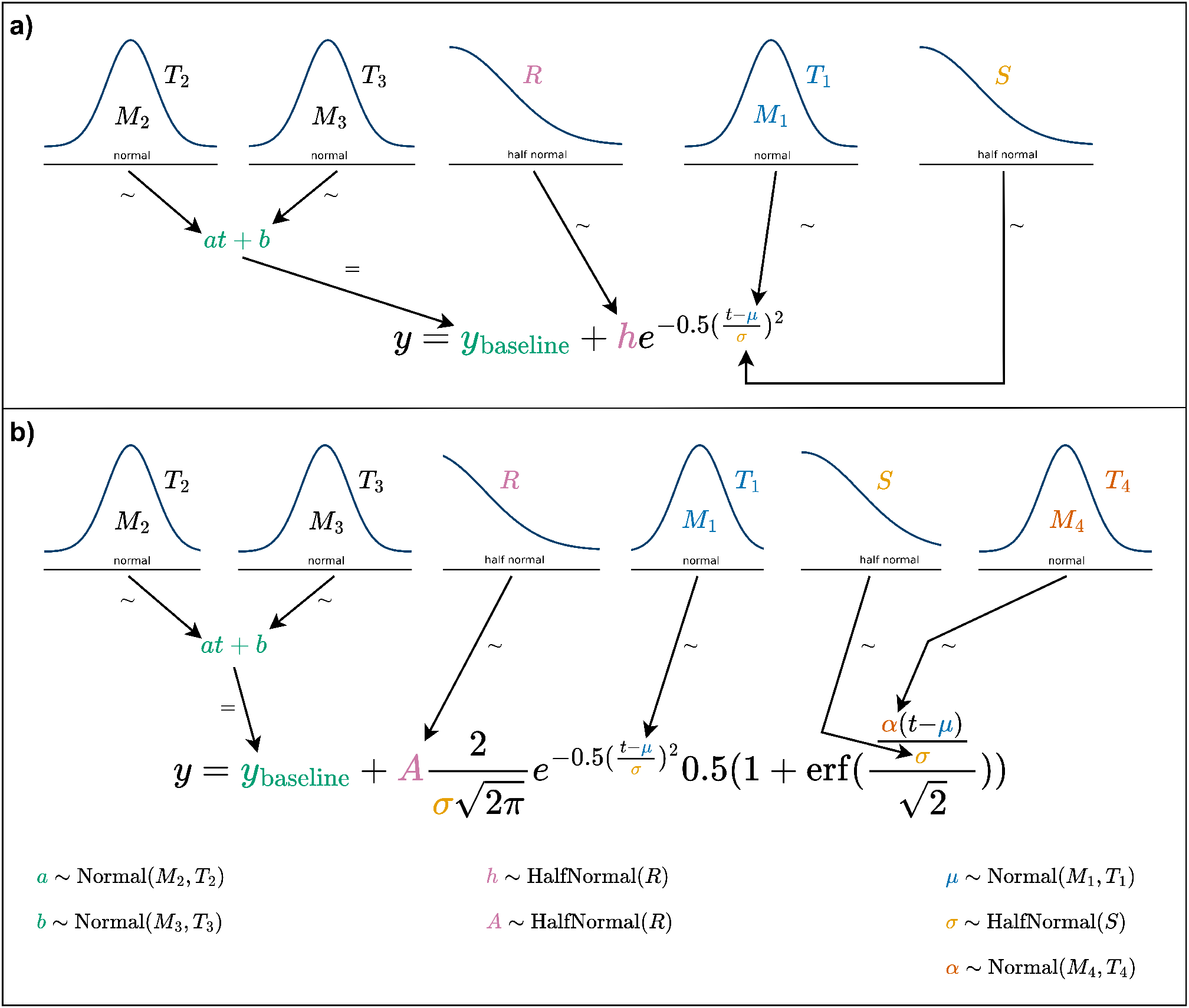
The intensity functions of normal (**a**) and skew normal peak models (**b**) as well as the prior probability distributions of their parameters are shown in the style of a Kruschke diagram [8]. Connections with ∼ imply stochastic and with = deterministic relationships. In case of variables with multiple occurrences in one formula, the prior was only connected to one such instance to preserve visual clarity. The variables *M*_*i*_ and *O*_*i*_ describe mean values and *T*_*i*_, *R*, and *S* standard deviations.

The double peak models (Figure 2) featured many of the same variables as their single peak counterparts so only the differences will be highlighted here. All variables pertaining to the actual peak were represented as vectors with two entries labeled with 0 and 1 by adding a named dimension to that effect. Aside from that, their priors remained unaltered except for the peak mean *µ*.

**Figure 2:**
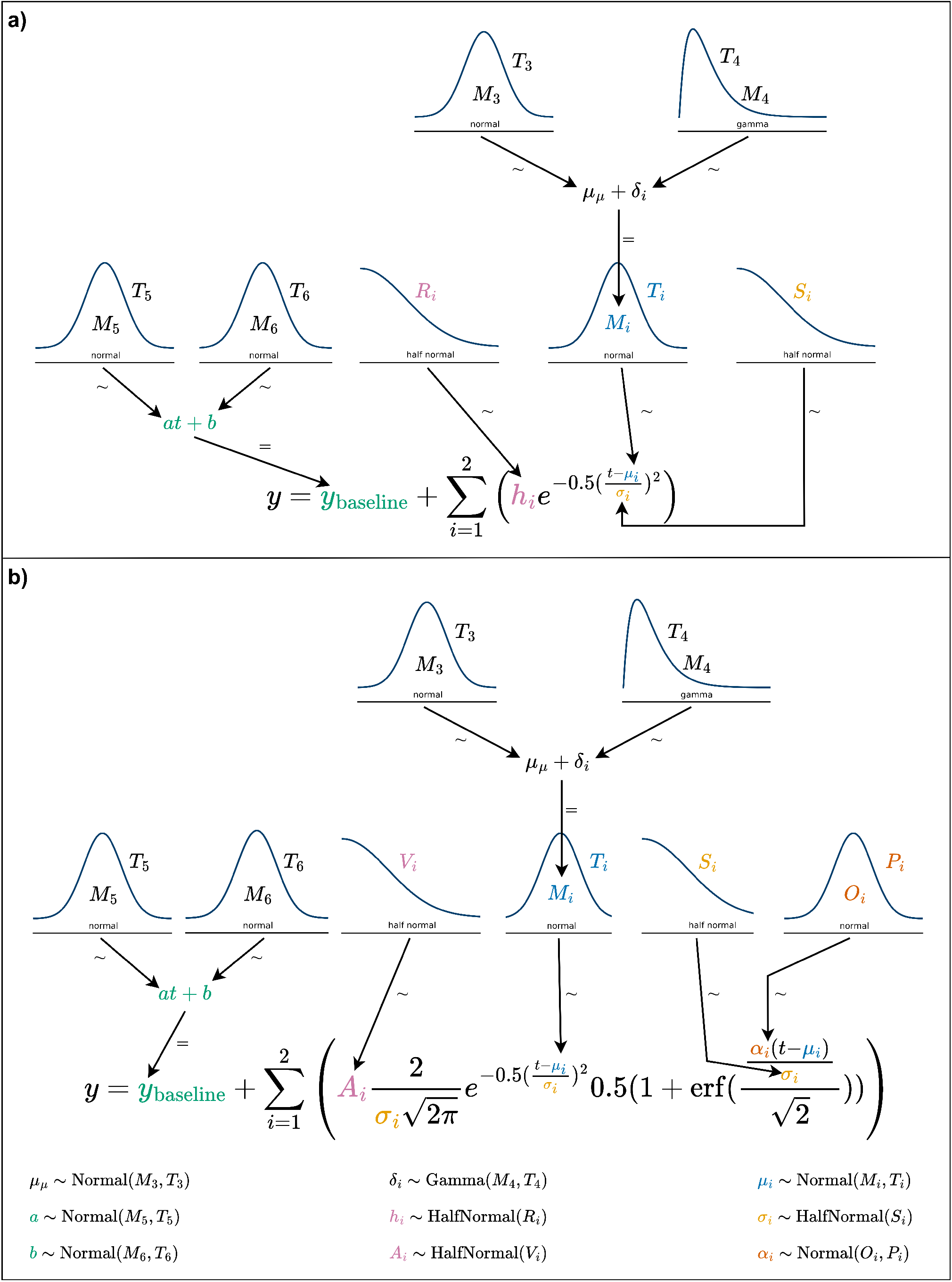
The intensity functions of double normal (**a**) and double skew normal peak models (**b**) as well as the prior probability distributions of their parameters are shown in the style of a Kruschke diagram [8]. Connections with ∼ imply stochastic and with = deterministic relationships. In case of variables with multiple occurrences in one formula, the prior was only connected to one such instance to preserve visual clarity. The variables *M*_*i*_ and *O*_*i*_ describe mean values and *T*_*i*_, *S*_*i*_, *P*_*i*_, and *V*_*i*_ standard deviations.

To provide a flexible solution to find double peak means across the whole time frame, the implementation of additional parameters proved indispensable. More precisely, the mean of both peaks or group mean was introduced as hyperprior (6) with a broad normal prior which enabled it to vary across the time frame as needed.

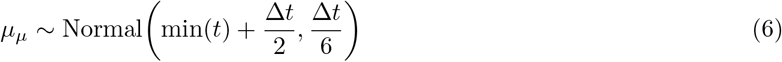

By defining a separation parameter representing the distance between the sub-peaks of a double peak

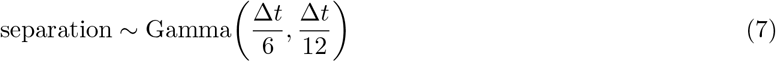

the offset of each peak’s mean parameter from the group mean is calculated as

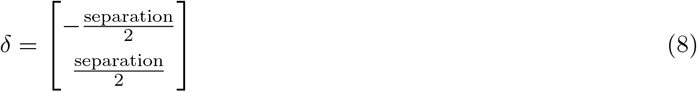

The priors for the mean parameters of each subpeak were then defined in dependence of *µ*_*µ*_ and δ as

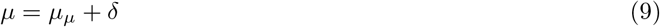

While all aforementioned parameters are necessary for the models, not all are of equal relevance for the user. A user’s primary interest for consecutive data analysis generally lies in obtaining mean values, peak areas and perhaps - usually to a much lesser degree - peak heights. Since only one of the latter two parameters is strictly required for scaling purposes, different models as shown in Figures 1 and 2 will feature either one or the other. Nonetheless, both peak area and peak height should be supplied to the user, hence the missing one was included as a deterministic model variable and thus still part of the model. In case of the normal and double normal models, the peak height *h* was used for scaling and the area *A* was calculated by

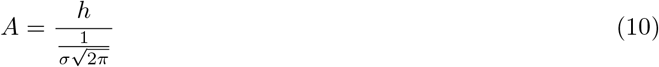

For skew normal and double skew normal models, the scaling parameter was the peak area. Since the mode and mean of a skewed distribution are – in contrast to normal distributions – distinct, the calculation of the height was nontrivial and ultimately a numerical approximation was added to the skewed models.

Beyond these key peak parameters, all PyMC models created by `PeakPerformance` contain additional constant data variables, and deterministic model variables. For example, the time series, i.e. the analyzed raw data, as well as the initial guesses for noise, baseline slope, and baseline intercept are kept as constant data variables to facilitate debugging and reproducibility. Examples for deterministic model variables in addition to peak area or height are the predicted intensity values and the signal-to-noise ratio defined here as

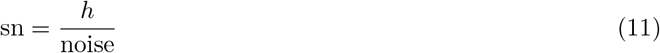

## 3. Results and Discussion

### 3.1 PeakPerformance workflow

`PeakPerformance` accommodates the use of a pre-manufactured data pipeline for standard applications as well as the creation of custom data pipelines using only its core functions. The provided data analysis pipeline was designed in a user-friendly way and requires minimal programming knowledge (Fig. 3). As portrayed in an example notebook in the code repository, only a few simple Python commands need to be executed. Instead of relying on these convenience functions, experienced users can also directly access the core functions of `PeakPerformance` for a more flexible application which is demonstrated in yet another example notebook. Before using `PeakPerformance`, the user has to supply raw data files containing a NumPy array with time in the first and intensity in the second dimension. For each peak, such a file has to be provided according to the naming convention specified in `PeakPerformance`’s documentation and gathered in one directory. If a complete time series of a 30 - 90 min LC-MS/MS run were to be submitted to the program, however, the target peak would make up an extremely small portion of this data. Additionally, other peaks with the same mass and fragmentation pattern may have been observed at different retention times. Therefore, it was decided from the outset that in order to enable proper peak fitting, only a fraction of such a time series with a range of 3 - 5 times the peak width and roughly centered on the target peak would be accepted as an input. This guarantees that there is a sufficient number of data points at the beginning and end of the time frame for estimating the baseline and noise level, as well.

**Figure 3:**
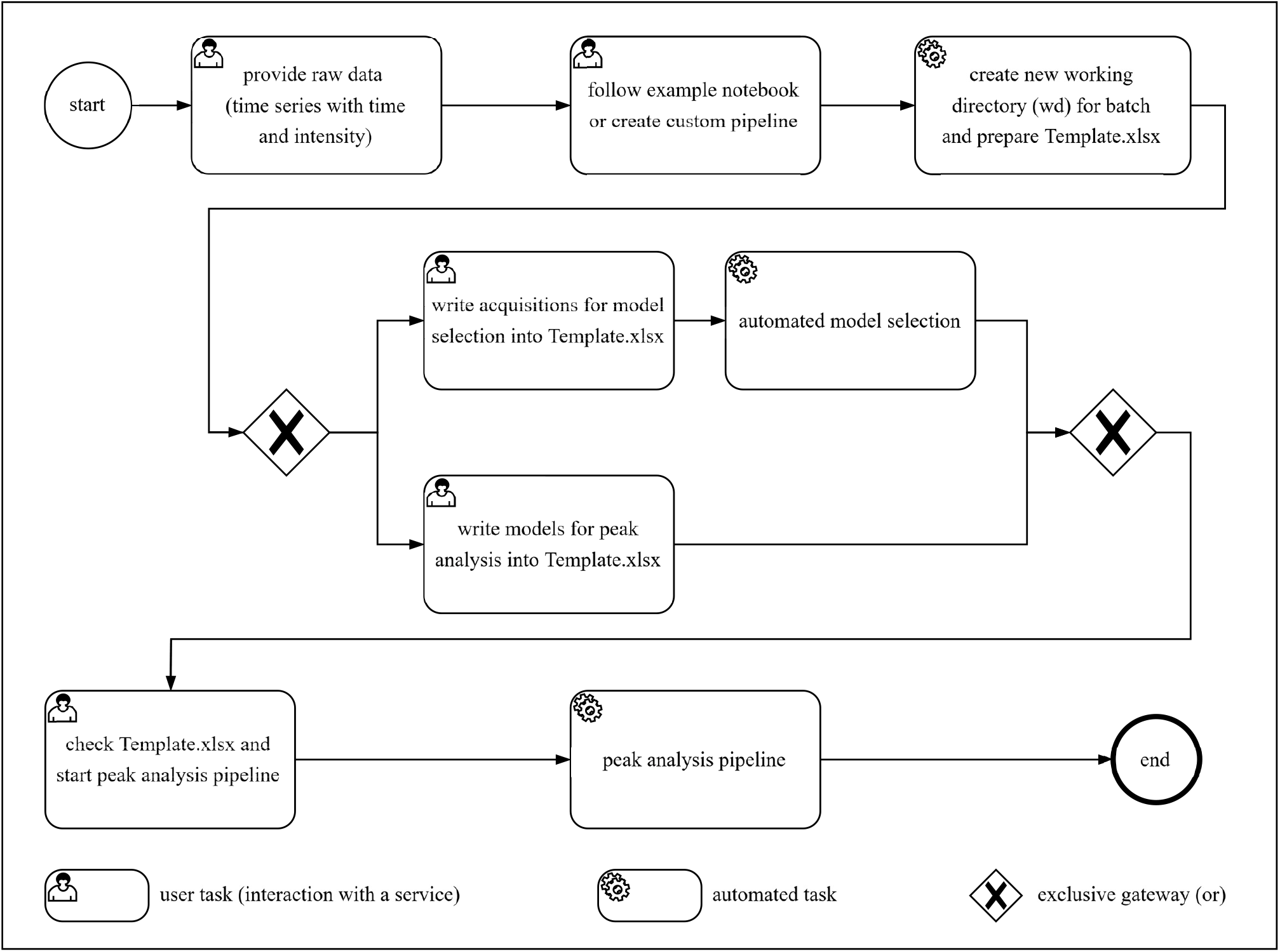
Overview of the pre-manufactured data analysis pipeline featured in `PeakPerformance`.

The provided data pipeline starts by defining a path to this raw data directory and one to a local clone of the `PeakPerformance` code repository. Using the `prepare model selection()` method, an Excel template file (“Template.xlsx”) for inputting user information is prepared and stored in the raw data directory. It is the user’s task, then, to select the settings for the pipeline within the file itself. Accordingly, the file contains detailed explanations of all parameters and the parsing functions of the software feature clear error messages in case mandatory entries are missing or filled out incorrectly.

Since targeted LC-MS/MS analyses essentially cycle through a list of mass traces for every sample, a model type has to be assigned to each mass trace. Preferably, this is done by the user which is of course only possible when the model choice is self-evident. If this is not the case, an optional automated model selection step can be performed, where one exemplary peak per mass trace is analyzed with all models to identify the most appropriate one. It is then assumed that within one batch run, all instances of a mass trace across all acquisitions can be fitted with the same type of model. For this purpose, the user must provide the name of an acquisition, i.e. sample, where a clear and representative peak for the given mass trace was observed. If e.g. a standard mixture containing all targets was measured, this would be considered a prime candidate. An additional feature lets the user exclude specific model types to save computation time and improve the accuracy of model selection by for example excluding double peak models when a single peak was observed. Upon provision of the required information, the automated model selection can be started using the `model selection()` function from the pipeline module and will be performed successively for each mass trace. Essentially, every type of model which has not been excluded by the user needs to be instantiated, sampled, and the log-likelihood needs to be calculated. Subsequently, the results for each model are ranked with the `compare()` function of the ArviZ package based on Pareto-smoothed importance sampling leave-one-out cross-validation (LOO-PIT) [9, 10]. This function returns a DataFrame showing the results of the models in order of their placement on the ranking which is decided by the expected log pointwise predictive density. The best model for each mass trace is then written to the Excel template file.

After a model was chosen either manually or automatically for each mass trace, the peak analysis pipeline can be started with the function `pipeline()` from the `pipeline` module. The first step consists of parsing the information from the Excel sheet. Since the pipeline, just like model selection, acts successively, a time series is read from its data file next and the information contained in the name of the file according to the naming convention is parsed. All this information is combined in an instance of `PeakPerformance`’s `UserInput` class acting as a centralized source of data for the program.

Depending on whether the “pre-filtering” option was selected, an optional filtering step will be executed to reject signals where clearly no peak is present before sampling, thus saving computation time. This filtering step uses the `find peaks()` function from the SciPy package [11] which simply checks for data points directly neighboured by points with lower intensity values. If no data points within a certain range around the expected retention time of an analyte fulfill this most basic requirement of a peak, the signal is rejected. Furthermore, if none of the candidate data points exceed a signal-to-noise ratio threshold defined by the user, the signal will also be discarded. Depending on the origin of the samples, this step may reject a great many signals before sampling saving potentially hours of computation time across a batch run of the `PeakPerformance` pipeline. For instance, in bioreactor cultivations, a product might be quantified but if it is only produced during the stationary growth phase, it will not show up in early samples. Another pertinent example of such a use case are isotopic labeling experiments for which every theoretically achievable mass isotopomer needs to be investigated, yet depending on the input labeling mixture, the majority of them might not be present in actuality.

Upon passing the first filter, a Markov chain Monte Carlo (MCMC) simulation is conducted using a No-U-Turn Sampler (NUTS) [12], preferably - if installed in the Python environment - the nutpie sampler [6] due to its highly increased performance compared to the default sampler of PyMC. Before sampling from the posterior distribution, a prior predictive check is performed the results of which can be accessed and evaluated after the fact.

When a posterior distribution has been obtained, the main filtering step is next in line. The first criterion is constituted by checking the convergence of the Markov chains towards a common solution for the posterior represented by the potential scale reduction factor [13], also referred to as the 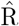 statistic or Gelman-Rubin diagnostic. If this factor is above 1.05 for any parameter, convergence was not reached and the sampling will be repeated once with a much higher number of tuning samples. If the filter is not passed a second time, the pertaining signal is rejected.

Harnessing the advantages of the uncertainty quantification, a second criterion calculates the ratio of the resulting standard deviation of a peak parameter to its mean and discards signals exceeding a threshold. Usually, false positives passing the first criterion are rather noisy signals where a fit was achieved but the uncertainty on the peak parameters is extremely high. These signals will then be rejected by the second criterion, ultimately reducing the number of false positive peaks significantly if not eliminating them.

If a signal was accepted as a peak, the final simulation step is a posterior predictive check which is added to the inference data object resulting from the model simulation.

### 3.2 Peak fitting results and diagnostic plots

After completing a cycle of the data pipeline or prematurely exiting it through one of the filters, the results need to be communicated and made available to the user. This is done in multiple ways. The most complete report is found in an Excel file called “peak data summary.xlsx”. Here, each analyzed time series has multiple rows (one per peak parameter) with the columns containing estimation results in the form of mean and standard deviation (sd) of the marginal posterior distribution, highest density interval (HDI), and the 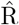 statistic among other metrics. Additional columns provide information on the acquisition and mass trace in question. Finally, there are columns stating whether the signal was recognized as a peak, if applicable the reason for the rejection of the signal, the utilized model for the simulation, and in case of a double peak a column specifying the peak number (“1st” or “2nd”). Accordingly, when a signal is rejected, it will nonetheless be added to the Excel report file and the exact reason for its rejection is detailed.

The second Excel file created is denominated as “area summary.xlsx” and is a more handy version of “peak data summary.xlsx” with a reduced degree of detail. As implied by the name, from the peak parameters only the peak area remains and the columns are trimmed down to the essentials. Since subsequent data analyses will most likely rely on the peak area, first and foremost, this sheet should facilitate the further usage of the data.

The most valuable result, however, is the storage of the inference data objects created only when a signal was identified as a peak. Conveniently, the inference data objects saved as `*.nc` files contain all data and metadata related to the process enabling the user to be able to perform diagnostics or create custom plots not featured in `PeakPerformance`.

Regarding data visualization with the matplotlib package [14, 15], `PeakPerformance`’s `plots` module offers the generation of two diagram types for each successfully fitted peak. The posterior plot presents the fit of the intensity function alongside the raw data points. The first row of Figure 4 presents two such examples where the single peak diagram shows the histidine (His) fragment with a m/z ratio of 110 Da and the double peak diagram the leucine (Leu) and isoleucine (Ile) fragments with a m/z ratio of 86 Da. The posterior predictive plots in the second row of Figure 4, then, exhibit the posterior predictive checks alongside the raw data. Since a posterior predictive check is based on drawing samples from the likelihood function, the result represents the theoretical range of values encompassed by the model. Accordingly, this plot enables users to judge whether the selected model can accurately explain the data.

**Figure 4:**
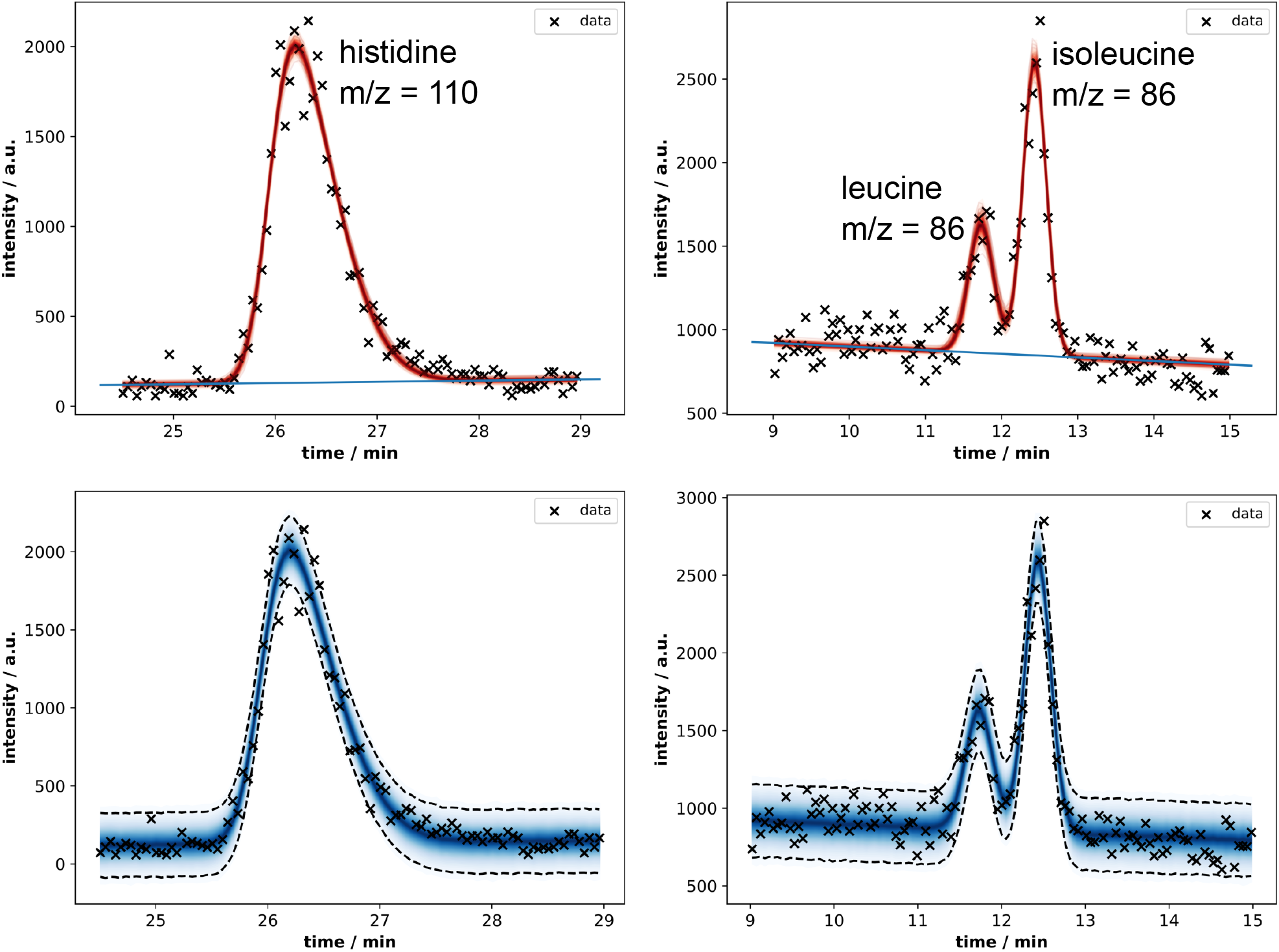
Results plots for a single His peak and a double Leu and Ile peak depicting the peak fit (first row) and the posterior predictive checks (second row) alongside the raw data. The numerical results are listed in table 2.

To complete the example, Table 2 shows the results of the fit in the form of mean, standard deviation, and HDI of each parameter’s marginal posterior. In this case, the fits were successful and convergence was reached for all parameters. Most notably and for the first time, the measurement noise was taken into account when determining the peak area as represented by its standard deviation and as can be observed in the posterior predictive plots where the noisy data points fall within the boundary of the 95 % HDI.

**Table 2:** Depiction of the results for the most important peak parameters of a single peak fit with the skew normal model and a double peak fit with the double normal model. Mean, area, and height have been highlighted in bold print as they constitute the most relevant parameters for further data evaluation purposes. The results correspond to the fits exhibited in Figure 4.

| parameter | single peak (skew normal distribution) |  |  |  | double peak (double normal distribution) |  |  |  |
| --- | --- | --- | --- | --- | --- | --- | --- | --- |
|  | mean | sd | hdi_3% | hdi_97% | mean | sd | hdi_3% | hdi_97% |
| baseline_intercept | -43.94 | 7.41 | -57.88 | -30.02 | 1115.40 | 38.69 | 1040.14 | 1185.07 |
| baseline_slope | 6.66 | 0.51 | 5.71 | 7.63 | -21.65 | 3.09 | -27.50 | -15.94 |
| noise | 103.63 | 7.51 | 89.50 | 117.26 | 118.63 | 8.01 | 103.52 | 133.29 |
| mean | **25.95** | **0.01** | 25.93 | 25.97 | **11.73** | **0.02** | 11.70 | 11.76 |
|  |  |  |  |  | **12.43** | **0.01** | 12.42 | 12.45 |
| area | **1512.32** | **37.31** | 1441.25 | 1581.37 | **317.16** | **28.84** | 263.23 | 370.56 |
|  |  |  |  |  | **674.34** | **26.34** | 623.47 | 722.88 |
| height | **1879.72** | **37.71** | 1809.30 | 1950.64 | **774.99** | **65.28** | 653.50 | 897.88 |
|  |  |  |  |  | **1762.66** | **64.04** | 1639.62 | 1881.66 |
| std | 0.53 | 0.02 | 0.48 | 0.56 | 0.16 | 0.02 | 0.13 | 0.20 |
|  |  |  |  |  | 0.15 | 0.01 | 0.14 | 0.17 |
| sn | 18.24 | 1.37 | 15.69 | 20.76 | 6.56 | 0.72 | 5.22 | 7.88 |
|  |  |  |  |  | 14.93 | 1.14 | 12.82 | 17.14 |
| alpha | 2.96 | 0.39 | 2.27 | 3.71 | - | - | - | - |

Another important feature of `PeakPerformance` is constituted by the easy access to diagnostic metrics for extensive quality control. Using the data stored in an inference data object of a fit, the user can utilize the ArviZ package to generate various diagnostic plots. One particularly useful one is the cumulative posterior predictive plot portrayed in Figure 5. This plot enables users to judge the quality of a fit and identify instances of lack-of-fit. As can be seen in the left plot, some predicted intensity values in the lowest quantile of the single peak example show a minimal lack-of-fit. Importantly, such a deviation can be observed, judged and is quantifiable which intrinsically represents a large improvement over the status quo.

**Figure 5:**
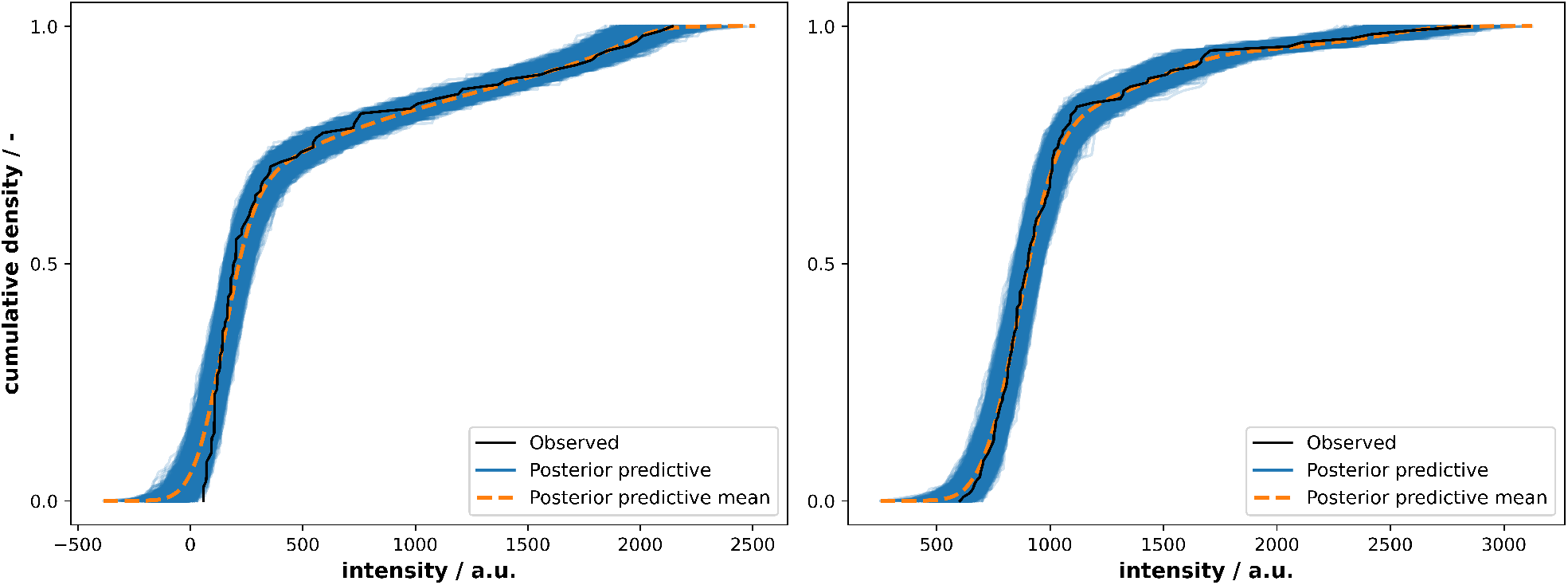
Cumulative posterior predictive plots created with the ArviZ package and pertaining to the example data of the single His peak (left) and the double Leu and Ile peak (right).

### 3.3 Validation

In the first stage of validation, peak fitting with normal and skew normal peak models was tested regarding the ability to reproduce the ground truth of randomly generated noisy synthetic data sets. The arithmetic means portrayed in Figure 6a were calculated based on a measure of similarity

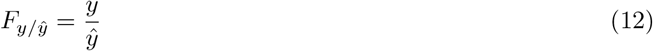

where *y* represents the estimated parameter value and 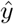 its pertaining ground truth. As they exhibit values close to 1, this demonstrates a near identity between estimation and ground truth. Additionally, the normal-shaped peak model was paired with skew normally distributed noisy data and vice versa. In both cases, *σ* was not reproduced well, especially by the normal-shaped model. Nevertheless, the peak area and height were still identified correctly with the skew normal model and merely slightly underestimated by the normal model. In the second stage, marginal cases in the form of slightly skewed peaks were investigated to observe whether their estimation with a normal- or skew normal-shaped intensity function would result in significant differences in terms of peak area and height. Here, a slight skew was defined as an *α* parameter of 1 resulting in peak shapes not visibly discernible as clearly normal or skew normal. With a sample size of 100 noisy, randomly generated data sets, we show that nearly identical estimates for peak area and height, as well as their respective uncertainties are obtained regardless of the utilized model (Fig. 6b). The exhibited mean values are based on fractions of the key peak parameters area and height between results obtained with a normal and skew normal model which were defined as

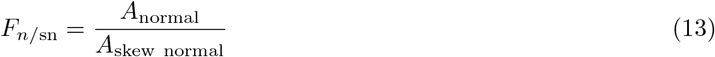

where *A*_*normal*_ and *A*_*skew normal*_ are the estimated areas with normal and skew normal models, respectively. In the third stage, experimental peak data was analyzed with both `PeakPerformance` (version 0.7.0) and Sciex MultiQuant (version 3.0.3) and the fraction of the obtained areas was determined as

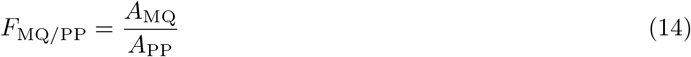

where *A*_MQ_ denominates the area yielded by MultiQuant and *A*_PP_ the area from `PeakPerformance`. Beyond the comparability of the resulting peak area ratio means portrayed in Figure 6c, it is relevant to state that 103 signals from MultiQuant (54 % of total signals) were manually modified. Of these, 31 % were false positives and 69 % were manually re-integrated. These figures are the result of a relatively high share of double peaks in the test sample which generally give a lot more cause for manual interference than single peaks. In contrast, however, the `PeakPerformance` pipeline was only started once and merely two single peaks and one double peak were fit again with a different model and/or increased sample size after the original pipeline batch run had finished. Among the 192 signals of the test data set, there were 7 noisy, low intensity signals without a clear peak which were recognized as a peak only by either one or the other software and were hence omitted from this comparison. By showing not only the mean area ratio of all peaks but also the ones for the single and double peak subgroups, it is evident that the variance is significantly higher for double peaks. In case of this data set, two low quality double peaks in particular inflated the variance significantly which may not be representative for other data sets. It has to be stated, too, that the prevalence of manual re-integration of double peaks in MQ might have introduced a user-specific bias, thereby increasing the final variance. Nevertheless, it could be shown that `PeakPerformance` yields comparable peak area results to a commercially available vendor software.

**Figure 6:**
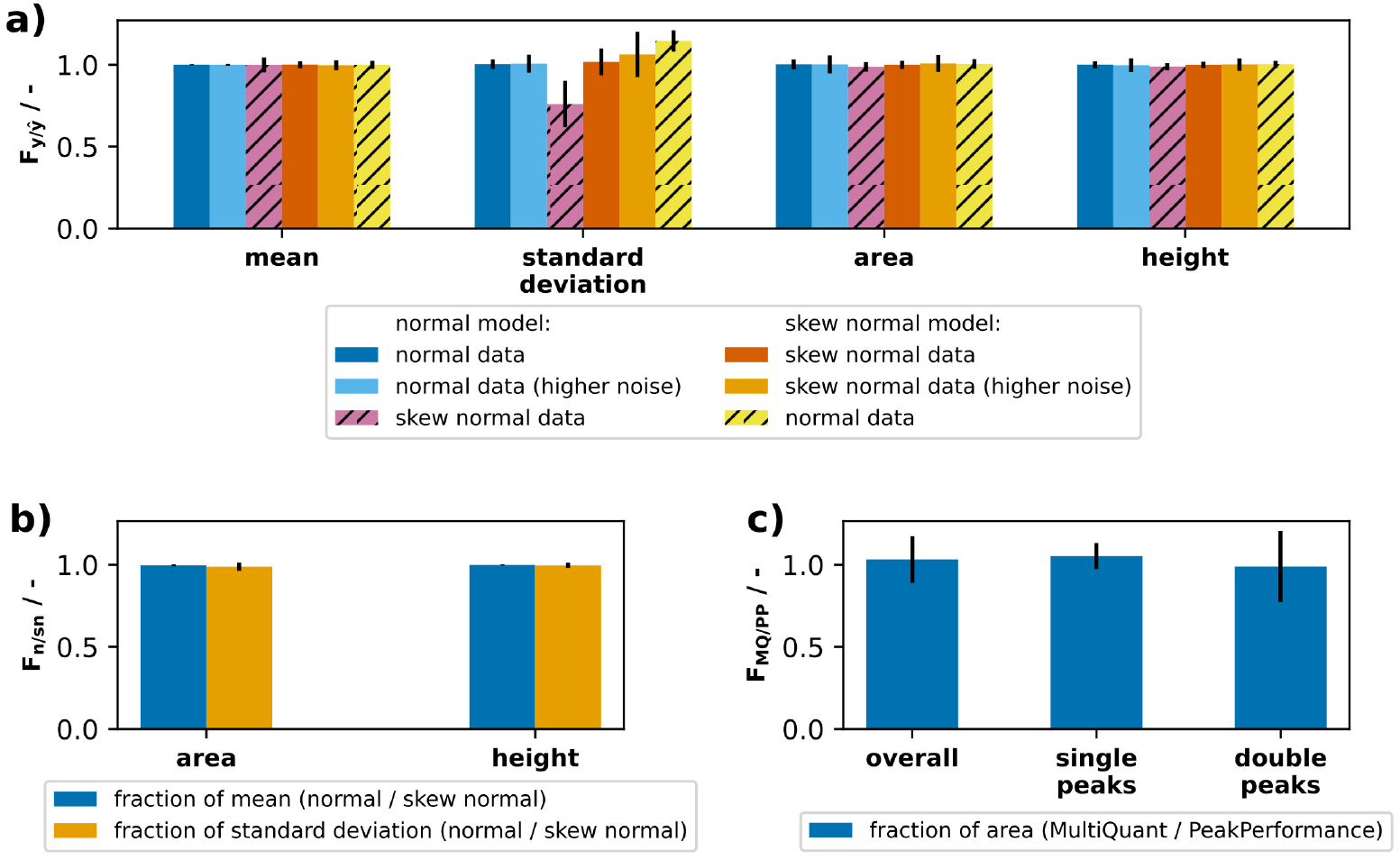
Validation of results from `PeakPerformance`. **a)** Noisy synthetic data was randomly generated from one of the implemented distributions and the program’s ability to infer the ground truth was observed. Portrayed are the fractions of estimated parameter to ground truth. **b)** The influence of model choice between normal and skew normal model in marginal cases with little to no skew was tested and the ratios between results from both models are plotted. **c)** Lastly, experimental data was analyzed with `PeakPerformance` version 0.7.0 and compared to results achieved with the commercial software Sciex MultiQuant version 3.0.3.

## 4 Conclusions

`PeakPerformance` is a tool for automated LC-MS/MS peak data analysis employing Bayesian inference. Due to its Bayesian methodology, it provides built-in uncertainty quantification and thus for the first time takes the measurement noise of an LC-MS/MS device into account when integrating peaks. Further, it allows strict control of model selection by the user but is not dependent on it due to the integrated automated model selection functionality. Regarding peak acceptance, it improves on vendor software solutions with more sophisticated, multi-layered metrics for decision making based on convergence of the parameter estimation, as well as the uncertainties of peak parameters. Finally, it allows the addition of new models to describe peak intensity functions with just a few minor code changes, thus lending itself to expansion to data from other chromatographic techniques. The design of `PeakPerformance` accommodates users with little programming experience by supplying convenience functions and relying on Microsoft Excel for data input and reporting. Its code repository on GitHub features automated unit tests, and the automatically built documentation (https://peak-performance.rtfd.io) includes instructions on how to apply `PeakPerformance` for your own chromatographic data which likely requires an adaptation of model priors. In future releases of `PeakPerformance`, we intend to implement an even more customizable building-block assembly of multi peak models, as well the extension of `PeakPerformance` to HPLC data.

## 5 Author contributions statement

`PeakPerformance` was conceptualized by JN and MO. Software implementation was conducted by JN with code review by MO. The original draft was written by JN with review and editing by MO, SN, EvL, and WW. The work was supervised by SN and funding was acquired by SN, EvL, and WW.

## 6 Acknowledgments

The authors thank Tobias Latour for providing experimental LC-MS/MS data for the comparison with the vendor software MultiQuant. Jochen Nießer and Michael Osthege would also like to express their gratitude towards Prof. Büchs, whose passion for empirical and kinetic modeling has inspired them and shaped their professional career and choice of methods. From your lectures we learned to embrace the richness of time-series data and the quantifiable insight that one can gain by the application of mathematical models.

## Abbreviations

HDI: highest density interval
His: histidine
Ile: isoleucine
LC-MS/MS: liquid chromatography tandem mass spectrometry
Leu: leucine
LOO-PIT: Pareto-smoothed importance sampling leave-one-out cross-validation
PDF: probability density function
prior: prior probability distribution
MCMC: Markov chain Monte Carlo
NUTS: No-U-Turn Sampler
sd: standard deviation
TMID: tandem mass isotopomer distribution

## References

[1] J. Hemmerich, S. Noack, W. Wiechert, and M. Oldiges. Microbioreactor systems for accelerated bioprocess development. Biotechnol J, 13(4):e1700141, 2018.

[2] Y. Kostov, P. Harms, L. Randers-Eichhorn, and G. Rao. Low-cost microbioreactor for high-throughput bioprocessing. Biotechnol Bioeng, 72(3):346–52, 2001.

[3] A. Azzalini. A class of distributions which includes the normal ones. Scand. J. Statist., 12:171–178, 1985.

[4] E. Grushka. Characterization of exponentially modified gaussian peaks in chromatography. Anal Chem, 44(11):1733–8, 1972.

[5] O. Abril-Pla, V. Andreani, C. Carroll, L. Dong, C. J. Fonnesbeck, M. Kochurov, R. Kumar, J. Lao, C. C. Luhmann, O. A. Martin, M. Osthege, R. Vieira, T. Wiecki, and R. Zinkov. PyMC: a modern, and comprehensive probabilistic programming framework in python. PeerJ Comput Sci, 9:e1516, 2023.

[6] Adrian Seyboldt and PyMC Developers. nutpie.

[7] Ravin Kumar, Colin Carroll, Ari Hartikainen, and Osvaldo Martin. Arviz a unified library for exploratory analysis of bayesian models in python. Journal of Open Source Software, 4(33), 2019.

[8] John K. Kruschke. Doing Bayesian Data Analysis. 1st edition edition, 2010.

[9] Sumio Watanabe. Asymptotic equivalence of bayes cross validation and widely applicable information criterion in singular learning theory. Journal of machine learning research, 11:3571–3594, 2010.

[10] Aki Vehtari, Andrew Gelman, and Jonah Gabry. Practical bayesian model evaluation using leave-one-out cross-validation and waic. Statistics and Computing, 27(5):1413–1432, 2016.

[11] Pauli Virtanen, Ralf Gommers, Travis E. Oliphant, Matt Haberland, Tyler Reddy, David Cournapeau, Evgeni Burovski, Pearu Peterson, Warren Weckesser, Jonathan Bright, Stéfan J. van der Walt, Matthew Brett, Joshua Wilson, K. Jarrod Millman, Nikolay Mayorov, Andrew R. J. Nelson, Eric Jones, Robert Kern, Eric Larson,J J Carey, İlhan Polat, Yu Feng, Eric W. Moore, Jake VanderPlas, Denis Laxalde, Josef Perktold, Robert Cimrman, Ian Henriksen, E. A. Quintero, Charles R. Harris, Anne M. Archibald, AntÄnio H. Ribeiro, Fabian Pedregosa, Paul van Mulbregt, and SciPy 1.0 Contributors. SciPy 1.0: Fundamental Algorithms for Scientific Computing in Python. Nature Methods, 17:261–272, 2020.

[12] Matthew D. Hoffmann and Andrew Gelman. The no-u-turn sampler: Adaptively setting path lengths in hamiltonian monte carlo. Journal of Machine Learning Research, 15, 2014.

[13] Andrew Gelman and Donald B. Rubin. Inference from iterative simulation using multiple sequences. Statistical Science, 7(4), 1992.

[14] J. D. Hunter. Matplotlib: A 2d graphics environment. Computing in Science & Engineering, 9(3):90–95, 2007.

[15] The Matplotlib Development Team. Matplotlib: Visualization with python, May 2024.

